# Small Directional Treadmill Perturbations Induce Differential Gait Stability Adaptation

**DOI:** 10.1101/2021.02.27.433210

**Authors:** Jinfeng Li, Helen J. Huang

## Abstract

Introducing unexpected perturbations to challenge gait stability is an effective approach to investigate balance control strategies. Little is known about the extent to which people can respond to small perturbations during walking. This study aimed to determine how subjects adapted gait stability to multidirectional perturbations with small magnitudes applied on a stride-by-stride basis. Ten healthy young subjects walked on a treadmill that either briefly decelerated belt speed (“stick”), accelerated belt speed (“slip”), or shifted the platform medial-laterally at right leg mid-stance. We quantified gait stability adaptation in both anterior-posterior and medial-lateral directions using margin of stability and its components, base of support and extrapolated center of mass. Gait stability was disrupted upon initially experiencing the small perturbations as margin of stability decreased in the stick, slip, and medial shift perturbations and increased in the lateral shift perturbation. Gait stability metrics were generally disrupted more for perturbations in the coincident direction. Subjects employed both feedback and feedforward strategies in response to the small perturbations, but mostly use feedback strategies during adaptation. Subjects primarily used base of support (foot placement) control in the lateral shift perturbation and extrapolated center of mass control in the slip and medial shift perturbations. These findings provide new knowledge about the extent of gait stability adaptation to small magnitude perturbations applied on a stride-by-stride basis and reveal potential new approaches for balance training interventions to target foot placement and center of mass control.

**NEW & NOTEWORTHY:** Little is known about if and how humans can adapt to small magnitude perturbations experienced on a stride-by-stride basis during walking. Here, we show that even small perturbations disrupted gait stability and that subjects could still adapt their reactive balance control. Depending on the perturbation direction, subjects might prefer adjusting their foot placement over their center of mass and vice versa. These findings could help potentially tune balance training to target specific aspects of balance.

## 1. Introduction

Perturbations during walking can induce potential losses of balance and elicit corrective locomotor adaptations, which are useful for understanding human balance control (1, 2). There are multiple approaches to perturb gait stability, including split-belt walking (1, 3), visual flow distortions (4, 5), waist pulling (6, 7), platform displacements (8, 9), and rapid changes in treadmill belt speed (10, 11). Studies using discrete perturbations such as waist pulling, platform displacements, and changes in treadmill belt speeds often apply the perturbation once out of every 5-20 strides (6–11). These large magnitude perturbations are less frequent and often require recovery steps to regain balance. Currently, there is no clear optimal frequency and magnitude for perturbation-based balance training during walking (12, 13), but more frequent exposure to perturbations seems to be beneficial for developing long-term fall-resisting skills (14, 15). Small magnitude perturbations are also more easily navigated by older adults (16, 17). As such, applying frequent repeated perturbations with small magnitudes during walking to create small but frequent losses of balance could be an effective approach for balance training during walking. Little is known, however, about the extent to which humans can adapt to small magnitude discrete balance perturbations applied on a stride-by-stride basis during walking. This approach could reveal stride-to-stride adaptation to almost imperceptible perturbations to gait stability, which would provide additional insight about how humans control, maintain, and adapt their stability during walking.

The margin of stability is a well-accepted metric to quantify stability of human walking (18). The margin of stability is the distance between the edge of the base of support and the extrapolated center of mass (19). People tend to maintain a constant margin of stability when walking on different surfaces (20, 21). When external conditions in the environment disrupt the steady state of walking and its margin of stability, increasing the base of support is effective for maintaining stability based on the inverted pendulum model (22) and has been demonstrated in multiple gait studies (6, 23, 24). For steps before and after unexpected treadmill belt acceleration perturbations during walking, one study showed that margin of stability decreased, and subjects took fewer recovery steps, suggesting improved or preserved stability (25). Treadmill belt speed induced backward loss-of-balance perturbations can lead to larger margin of stability reductions in older adults compared to young adults (26). Interestingly, these margin of stability responses scaled with the perturbation intensity in both the young and older adults (26). The extent to which humans can control their margin of stability in response to small magnitude perturbations applied on a stride-by-stride basis remains unknown.

Analyzing the base of support and the extrapolated center of mass components of margin of stability could provide insights on how the body reacts to perturbations and maintains stability. Controlling foot placement through the swing leg or controlling center of mass through the stance leg are two main balance strategies for perturbed walking (6, 27, 28). In young adults, adaptation to multi-directional waist pulling perturbations have resulted in increased medial-lateral foot placement at perturbation onset, potentially to proactively regulate balance and also increased anterior-posterior foot placement and margin of stability at contralateral heel strike, perhaps to reactively regain stability (29). Older adults have demonstrated that they tend to adjust their extrapolated center of mass, potentially to transfer the improved stability of the trained leg to the untrained leg in response to treadmill belt acceleration perturbations (30). Treadmill belt slip perturbations have also resulted in the lengthening of the base of support and shortening the forward excursion of the extrapolated center of mass in post-stroke individuals, effectively increasing their anterior-posterior margin of stability and overall stability (31). Thus, separating margin of stability into its components, the base of support and extrapolated center of mass, will provide greater insight on how humans adapt, if at all, to frequent small magnitude perturbations.

The primary purpose of this study was to determine how small treadmill perturbations applied on a stride-by-stride basis in healthy young adults affected margin of stability and its components (base of support and extrapolated center of mass). We rapidly accelerated or decelerated treadmill belt speed to create “slip” or “stick” perturbations, respectively. We also rapidly shifted (i.e., translated) the treadmill medially or laterally to create treadmill shift perturbations in the medial or lateral direction, respectively. We applied the treadmill perturbations at right mid-stance of each stride, except for no-perturbation “catch” strides that occurred randomly in each batch of 5 strides. The purpose of the “catch” strides was to probe whether subjects were potentially adopting a feedforward strategy in response to the small magnitude perturbations encountered during perturbed walking. Experimental protocols that apply perturbations trial-by-trial with unexpected catch trials have typically been used in goal-directed arm reaching studies (32, 33). We are extending that protocol to walking to gain insights on the adaptability of balance control during walking. Because we applied perturbations on a stride-by-stride basis, the strengths of the perturbations were limited to avoid the need for recovery steps, which are often necessary during larger treadmill induced trip and slip perturbations that seek to induce a fall.

The first hypothesis was that when subjects first experienced the small perturbations, their gait stability would be disrupted (i.e., become less stable) and then adapt to regain stability as they gained more experience with the perturbations. As such, we expected margin of stability to decrease upon first experiencing the perturbations and then increase over time with more exposure to the perturbations. The second hypothesis was that subjects would adopt more feedforward strategies as they gained more experience with the perturbations. We expected that margin of stability, base of support, and extrapolated center of mass during catch strides would differ from the averaged levels of the unperturbed strides prior to the perturbed period and deviate more as subjects gained more experience with the perturbations.

To assess the extent of the expected adaptation, we also sought to determine whether perturbation direction and size affected the responses and adaptation to the perturbations. We hypothesized that the anterior-posterior margin of stability and its components would be more sensitive to the treadmill belt perturbations, which were in the coincident (anterior-posterior) direction and likewise, medial-lateral margin of stability and its components would be more sensitive to the treadmill shift perturbations, which were in the coincident (medial-lateral) direction. We also hypothesized that adaptation of margin of stability and its components would scale with perturbation size.

## 2. Methods

### 2.1. Participants, setup and protocol

Ten healthy young subjects (4 males, 6 females; age: 21.7 ± 2.4 years) participated in this study. The Institutional Review Board of the University of Central Florida approved the experimental protocol (SBE-16-12831). The study met all requirements in accordance with the Declaration of Helsinki, and all subjects provided their written informed consent.

Subjects walked on a dual split-belt instrumented treadmill (M-gait, Motekforce Link, Amsterdam, The Netherlands) operating at 300 Hz as we recorded their lower limb kinematics using the 16-marker Conventional Lower Body Markerset and a passive motion capture system (OptiTrack, NaturalPoint Inc., Corvallis, OR, USA, 13 Prime13W and 9 Prime13 cameras) operating at 240 Hz. The treadmill has independent speed control for each treadmill belt, and the treadmill is mounted on a moveable platform such that the treadmill can rapidly translate side-to-side in real-time. These features were used to produce treadmill perturbations.

We programmed the treadmill (D-Flow, version 3.28.0, Motekforce Link, Amsterdam, The Netherlands) to apply perturbations during mid-stance of the right leg on a stride-by-stride basis. The perturbation onset occurred when the vertical projection of the estimated center of mass position (the average position of the four pelvis markers: left anterior superior iliac spine, right anterior superior iliac spine, left posterior superior iliac spine, and right posterior superior iliac spine) resided between the right toe (2^nd^ metatarsal) and right heel markers in the anterior-posterior direction and when the left heel marker was at least 5 cm higher than the right toe marker (Fig. 1A). The duration of each perturbation from onset to termination was ~400 ms such that the treadmill could return to the neutral unperturbed position and baseline belt speed (1.0 m/s) prior to or shortly after left heel strike.

**Fig. 1.**
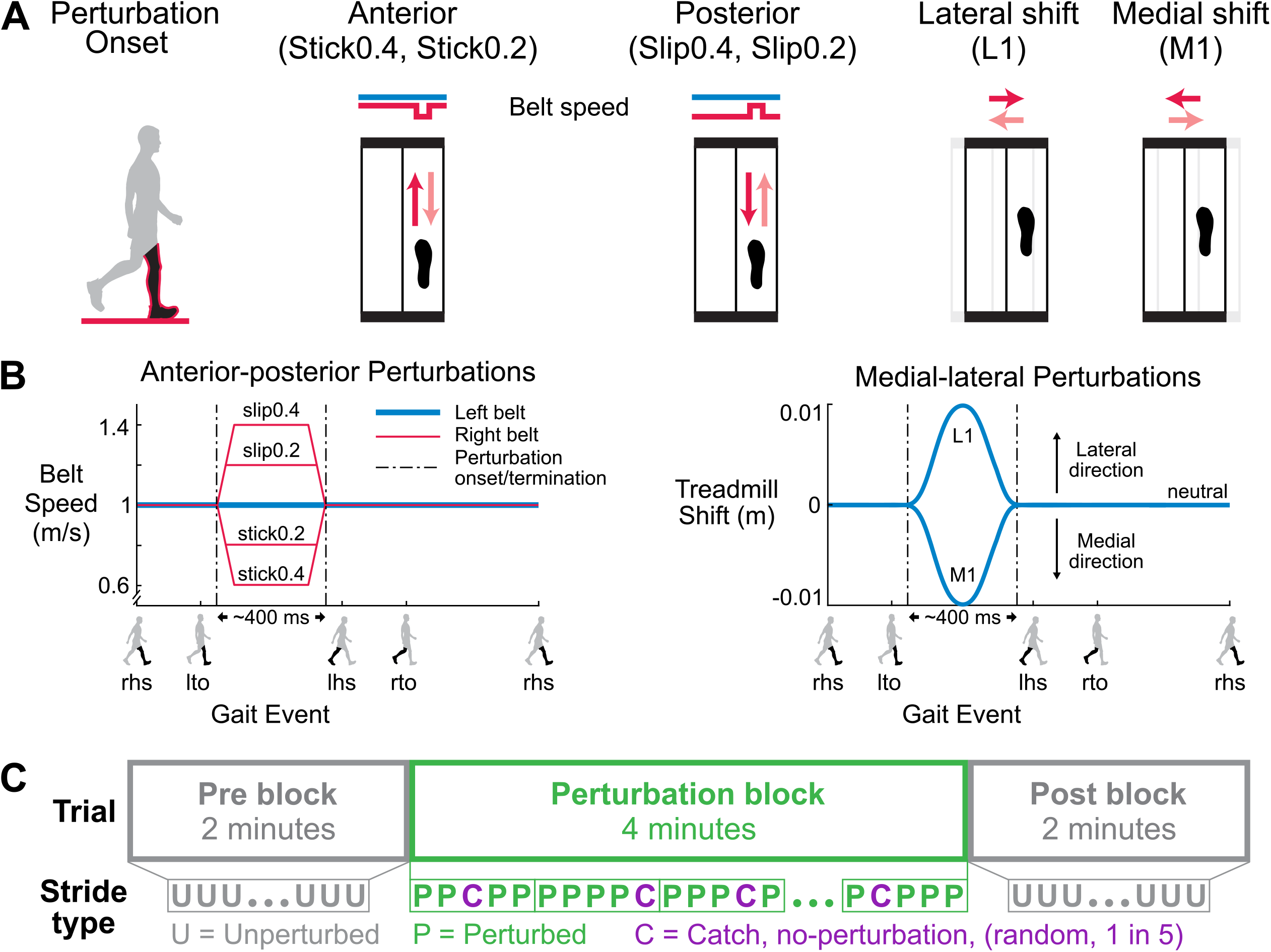
Schematic of the perturbations and experimental protocol. **(A)** Perturbation onset occurred at right leg mid-stance (black leg in red outline). There were 4 perturbation directions, and 2 sizes for the anterior-posterior perturbations, for a total of six perturbation conditions. The red arrows are the initial relative surface displacements, and the faded red arrows are the displacements to return to the unperturbed state. **(B)** The left belt speed was fixed at 1.0 m/s (blue). For the Stick0.2 and Stick0.4 perturbations, the right belt speed (red) decelerated to 0.8 m/s and 0.6 m/s, respectively, and then returned to the tied belt speed. For the Slip0.2 and Slip0.4 perturbations, the right belt speed accelerated to 1.2 m/s and 1.4 m/s, respectively, and then returned to the tied belt speed. For the L1 and M1 perturbations, the treadmill shifted 1 cm laterally and medially, respectively, and then returned to neutral. The perturbations were ~400 ms in duration. Gait events: right heel strike (rhs), left toe off (lto), left heel strike (lhs), and right toe off (rto). **(C)** Blocks of the experimental protocol. Each trial started with a 2-minute unperturbed walking block (pre), followed by a 4-minute perturbed walking block (perturbation), and completed with another 2-minute unperturbed walking block (post). An unperturbed catch stride occurred randomly 1 out of every 5 strides during the perturbation block.

Perturbations were applied in four directions: anterior, posterior, medial, and lateral to examine the effects of perturbation directions (Fig. 1A). We also used two perturbation sizes for the anterior and posterior directions, which could be completed in ~400 ms, to examine potential scaling effects with perturbation size (Fig. 1B). The baseline walking speed was 1.0 m/s, and the belt acceleration was set to 12.5 m/s^2^. For anterior “stick” perturbations, the right belt speed decreased from 1.0 m/s to the target speed (0.6 m/s and 0.8 m/s; Stick0.4 and Stick0.2, respectively) and then returned to the baseline speed. For posterior “slip” perturbations, the right belt speed increased to the target speed (1.2 m/s and 1.4 m/s; Slip0.2 and Slip0.4, respectively) and then returned to the baseline speed. For medial perturbations, the treadmill shifted 1 cm toward the medial side of the right leg (i.e., to the left) and then returned to the neutral treadmill location (M1). For lateral perturbations, the treadmill shifted 1 cm toward the lateral side of the right leg (i.e., to the right) and then returned to the neutral treadmill location (L1). The maximum acceleration for the medial-lateral perturbations was 3.6 m/s^2^.

Each subject completed six trials, one trial for each perturbation condition. We randomized the order of the six trials and did not inform subjects of the perturbation type, timing, direction, and magnitude. Each trial was 8 minutes long and consisted of 3 blocks. The trial began with 2 minutes of unperturbed walking (pre block), followed by 4 minutes of walking with perturbations (perturbation block) and concluded the trial with another 2 minutes of unperturbed walking (post block). During the 4-minute perturbation block, no-perturbation “catch” strides occurred randomly 1 out of every 5 strides to assess whether subjects were anticipating or reacting (Fig. 1C). Subjects wore a safety harness to prevent potential falls during all walking trials.

### 2.2. Data analysis

We wrote custom MATLAB (version 9.3, R2017b, Mathworks, Natick, MA, USA) scripts to process the kinematic data and calculate the gait stability metrics. We low-pass filtered the three-dimensional coordinates of the markers using a 4th order Butterworth filter with a 12 Hz cut-off frequency, and then extracted the gait events. Heel strike was the instant the vertical position of the heel marker of the foot transitioning from swing to stance was near zero (close to the treadmill) and reached its maximum vertical acceleration (34). Toe off was the instant the toe marker of the foot transitioning from stance to swing reached its maximum vertical acceleration between two consecutive heel strikes (35).

We calculated the margin of stability at every left heel strike (lhs) to assess reactive control and left toe off (lto) to assess anticipatory control as:

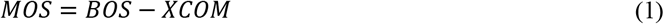

where MOS is the margin of stability; BOS is the boundary of the base of support; and XCOM is the extrapolated center of mass (19). We calculated the extrapolated center of mass using Equation 2, which also accounts for the movement velocity of the treadmill surface (36, 37).

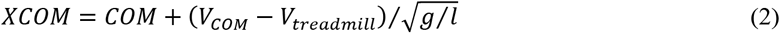

COM is the center of mass position, and V_COM_ is the center of mass velocity, which was derived as the first derivative of the center of mass position. V_treadmill_ is the belt velocity for the anterior-posterior perturbations and the platform velocity for the medial-lateral perturbations, respectively. The remaining variables are g, the gravitational constant (9.81 m/s^2^) and l, the equivalent pendulum length, which was the distance from the ankle marker (lateral malleolus) of the leading leg to the center of mass at left heel strike and left toe off.

We calculated the anterior-posterior margin of stability (MOS_ap_) as the anterior-posterior distance between the extrapolated center of mass and the toe marker of the leading leg in the sagittal plane, which defined the anterior-posterior base of support (BOS_ap_) (Fig. 2A). The anterior-posterior extrapolated center of mass (XCOM_ap_) was the anterior-posterior distance between the extrapolated center of mass and the toe marker of the trailing leg. We calculated the medial-lateral margin of stability (MOS_ml_) as the medial-lateral distance between the extrapolated center of mass and the ankle marker of the leading leg in the frontal plane, which defined the medial-lateral base of support (BOS_ml_) (Fig. 2B). The medial-lateral extrapolated center of mass (XCOM_ml_) was the medial-lateral distance between the extrapolated center of mass and the ankle marker of the trailing leg.

**Fig. 2.**
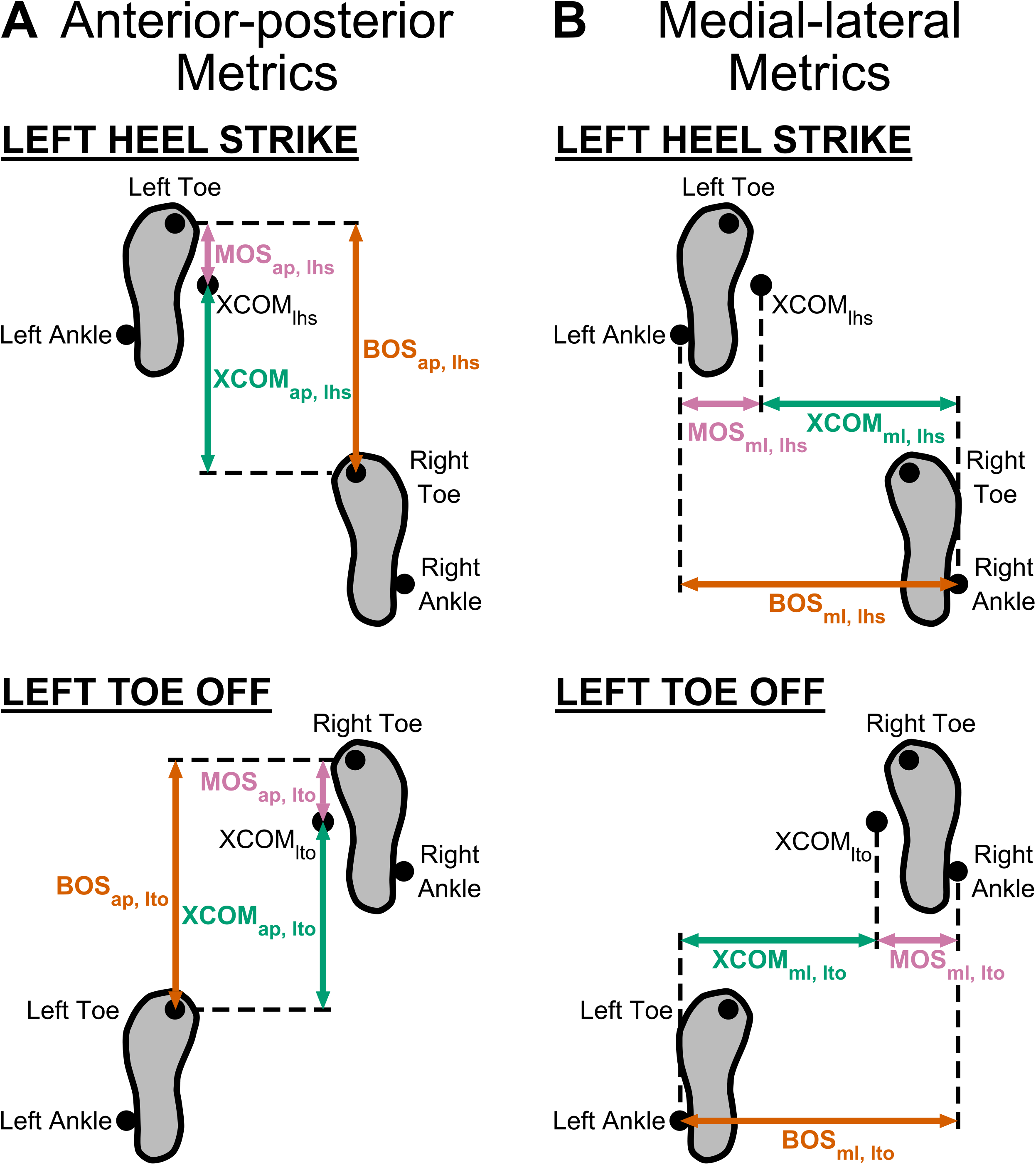
Anterior-posterior and medial-lateral metrics of gait stability at left heel strike and left toe off. **(A)** The anterior-posterior base of support (BOS_ap_, orange arrow) was the anterior-posterior distance of the toe marker of the leading leg relative to the toe marker of the trailing leg. The anterior-posterior extrapolated center of mass (XCOM_ap_, green arrow) was the anterior-posterior distance of XCOM relative to the toe marker of the trailing leg. The anterior-posterior margin of stability (MOS_ap_, pink arrow) was the BOS_ap_ - XCOM_ap_. **(B)** The medial-lateral base of support (BOS_ml_) was the medial-lateral distance of the ankle marker of the leading leg relative to the ankle marker of the trailing leg. The medial-lateral extrapolated center of mass (XCOM_ml_) was the medial-lateral distance of XCOM relative to the ankle marker of the trailing leg. The medial-lateral margin of stability (MOS_ml_) was the BOS_ml_ - XCOM_ml_.

We quantified margin of stability and its components at different phases throughout the protocol: pre, early perturbed, late perturbed, early catch, late catch, early post, and late post. Pre referred to all strides during the first 2-minute pre walking block except for the first 11 strides during which the treadmill belts were getting up to speed (38). Early perturbed was the first 3 perturbed strides during the 4-minute perturbation block. We used the first 3 perturbed strides as people tend to adapt quickly in arm reaching (39) and split-belt walking studies (40). For left toe off metrics, early perturbed used perturbed strides 2-4, which excluded the first perturbed stride since left toe off occurred before the perturbation onset. Late perturbed was the last 3 perturbed strides during the perturbation block, regardless of when the metrics were calculated. Early catch and late catch referred to the first 2 and last 2 unperturbed strides during the 4-minute perturbation block, respectively. Early post was the first 3 strides while late post was the last 3 strides during the last 2-minute post walking block.

### 2.3. Statistics

We performed statistical procedures using JMP (JMP Pro 12, SAS Institute Inc., Cary, NC, USA) and set the significance level to 0.05. First, we checked and excluded outliers as being beyond 3 standard deviations from the mean for each data set. We also used a Shapiro-Wilk W test, Levene’s test, and Mauchly’s test to check for the distribution normality, homogeneity of variance, and sphericity, respectively. For the first three hypotheses related to adaptation, using a feedforward strategy, and effects of perturbation directions, we analyzed the anterior (Stick0.4), posterior (Slip0.4), medial (M1), and lateral (L1) perturbation conditions. For the fourth hypothesis related to perturbation size effect, we analyzed the data of the 2 sizes of the anterior (Stick0.4, Stick0.2) and posterior (Slip0.4, Slip0.2) perturbations.

For the first hypothesis regarding adaptation, we performed one-way repeated measures analysis of variance (rANOVA) for the perturbed strides on each metric (MOS_ap, lhs_, MOS_ml, lhs_, MOS_ap, lto_, MOS_ml, lto_, BOS_ap, lhs_, BOS_ml, lhs_, BOS_ap, lto_, BOS_ml, lto_, XCOM_ap, lhs_, XCOM_ml, lhs_, XCOM_ap, lto_, and XCOM_ml, lto_) with phase (pre, early perturbed, late perturbed, early post, and late post) as a factor by perturbation condition. When phase was a main effect, we performed post hoc pairwise comparisons using Tukey’s Honest Significant Difference (HSD) tests to account for multiple comparisons for the following apriori comparisons: 1) pre to early perturbed to assess the disruption of gait stability, 2) early perturbed to late perturbed to assess adaptation, 3) early post to late post to assess de-adaptation. To determine which component of margin of stability contributed more to changes in margin of stability (|ΔMOS|), we compared the magnitudes of the changes of the base of support (|ΔBOS|) and extrapolated center of mass position (|ΔXCOM|) within a phase comparison (pre and early perturbed; early and late perturbed) using paired t-tests (i.e., for pre and early perturbed, |ΔBOS| = |BOS_pre_ − BOS_early_perturbed_| versus |ΔXCOM| = |XCOM_pre_ − XCOM_early_perturbed_|; for early to late perturbed, |ΔBOS| = |BOS_early_perturbed_ − BOS_late_perturbed_| versus |ΔXCOM| = |XCOM_early_perturbed_ − XCOM_late_perturbed_|).

For the second hypothesis regarding whether subjects used a feedforward strategy for adaptation, we performed one-way rANOVA for the catch strides on each metric with phase (pre, early catch, late catch) as a factor by perturbation condition. If phase was a main effect, we performed Tukey’s HSD post hoc tests for the following comparisons: 1) pre to early catch to assess anticipatory responses at early catch, 2) pre to late catch to assess anticipatory responses at late catch, and 3) early catch to late catch to assess if subjects used more feedforward strategies during adaptation.

For the third hypothesis regarding perturbation direction, we also used a one-way rANOVA for each metric (ΔMOS, ΔBOS, and ΔXCOM for early perturbed minus pre; ΔMOS, ΔBOS, and ΔXCOM for late perturbed minus early perturbed) with perturbation condition as a factor. Here, the direction of the change, not just the magnitude, from pre to early and from early to late was important. We performed post hoc pairwise comparisons using Tukey’s HSD if perturbation condition was a main effect.

For the fourth hypothesis regarding a scaling effect, we used a two-way rANOVA on each metric (|ΔMOS|, |ΔBOS|, and |ΔXCOM| for pre to early perturbed; |ΔMOS|, |ΔBOS|, and |ΔXCOM| for early to late perturbed) with perturbation condition (stick, slip) and size (regular, 0.4; half-size, 0.2) as factors. If a significant interaction effect was detected, we performed paired t-tests with Bonferroni correction (α = 0.025) to identify differences between sizes for each perturbation condition.

We calculated effect size using partial eta squared (η_p_^2^) for the rANOVA results and Cohen’s d for post hoc pairwise comparisons and paired t-tests (41). Complete statistical results are provided in the supplemental data (Supplemental Tables S1-S6; https://doi.org/10.6084/m9.figshare.16910683).

## 3. Results

### 3.1. Representative data

Changes of MOS and its components during perturbed walking were evident throughout the gait cycle (Fig. 3). When perturbations first occurred at early perturbed, MOS_lhs_, BOS_lhs_, and XCOM_lhs_ all deviated from the pre unperturbed strides, indicating that the perturbations disrupted the subjects. As subjects gained more experience with the perturbations, MOS_lhs_ and BOS_lhs_ of late perturbed generally trended back to pre levels, suggesting adaptation occurred. When the perturbations were removed at the post block, MOS_lhs_ and its components were almost the same as pre, suggesting a return to pre levels. The MOS_ap, lto_ of the Stick0.4 perturbation, BOS_ap, lto_ and XCOM_ap, lto_ of the Slip0.4 perturbation, and BOS_ml, lto_ and XCOM_ml, lto_ of the L1 perturbation deviated from pre at early perturbed, suggesting an anticipatory response. The BOS_lto_ and XCOM_lto_ of the Slip0.4 and L1 perturbations then trended to their pre levels by late perturbed, an indication of adaptation, and returned to the pre levels by the end of post.

**Fig. 3.**
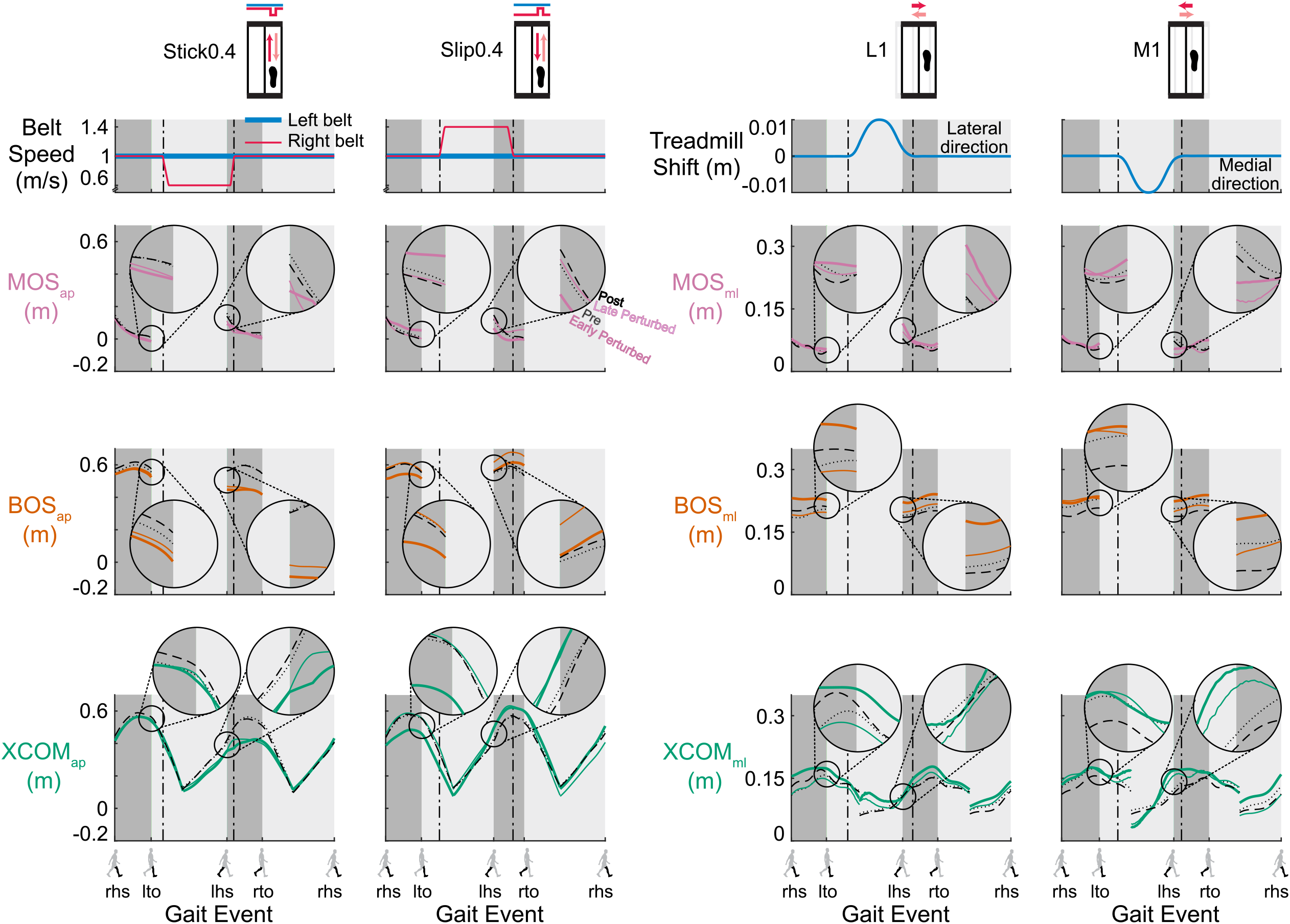
Trajectories of the gait stability metrics (MOS, pink; BOS, orange, and XCOM, green) for specific strides of a representative subject for the Stick0.4, Slip0.4, L1, and M1 perturbations (black dotted line: the 2^nd^ to last stride of pre; colored thick line: the 2^nd^ stride of early perturbed; colored thin line: the 2^nd^ to last stride of late perturbed; and black dashed line: the 2^nd^ to last stride of post). The MOS_ap, lhs_, BOS_ap, lhs_, and XCOM_ap, lhs_ of the Stick0.4 and Slip0.4 perturbations, and MOS_ml, lhs_, BOS_ml,_ lhs, and XCOM_ml, lhs_ of the L1 and M1 perturbations deviated from pre at early perturbed, then MOS_lhs_ and BOS_lhs_ generally trended to the pre trajectory by late perturbed and returned to the pre trajectory by post, shown in the exploded view in the inset circles. The MOS_ap, lto_ of the Stick0.4 perturbation, BOS_ap, lto_ and XCOM_ap, lto_ of the Slip0.4 perturbation, and BOS_ml, lto_ and XCOM_ml, lto_ of the L1 perturbation deviated from pre at early perturbed, then BOS_lto_ and XCOM_lto_ trended to the pre trajectory by late perturbed and returned to the pre trajectory by post, shown in the exploded view in the inset circles. Gait events: right heel strike (rhs), left toe off (lto), left heel strike (lhs), and right toe off (rto). Dark grey area: double support phase. Light grey area: single support phase. The plots show the MOS and BOS during double support and XCOM for the entire gait cycle.

### 3.2. Adaptation to small anterior-posterior perturbations applied on a stride-by-stride basis (hypothesis 1)

Subjects adapted to the Stick0.4 perturbation after an initial disruption to gait stability (Fig. 4A and Supplemental Fig. S1A, all supplemental figures are available at https://doi.org/10.6084/m9.figshare.16910713). Phase had a significant effect on all metrics except for BOS_ap, lto_ for the Stick0.4 perturbation (see Supplemental Table S1 for metric specific F, p, and η_p_^2^). Upon initial exposure to the perturbations, MOS_ap, lhs_ was disrupted and decreased significantly by 41.3% (p < 0.001, Cohen’s d = 2.49). After the initial perturbation, subjects immediately took shorter and wider steps and shifted their XCOM backward at left heel strike (pre to early perturbed: BOS_ap, lhs_ decreased, p < 0.001, Cohen’s d = 4.95; BOS_ml, lhs_ increased, p = 0.03, Cohen’s d = 0.99; XCOM_ap, lhs_ decreased, p < 0.001, Cohen’s d = 2.32). Starting with the second Stick0.4 perturbation, subjects also took wider steps (determined by the previous right heel strike) and shifted their XCOM forward at left toe off in anticipation of an impending perturbation at right leg mid-stance (pre to early perturbed: BOS_ml, lto_ increased, p < 0.001, Cohen’s d = 2.21; XCOM_ap, lto_ increased, p < 0.001, Cohen’s d = 1.83). MOS_ap, lhs_ increased significantly by 44.4% (p < 0.001, Cohen’s d = 1.57) after the initial disruption and trended toward pre level by late perturbed, indicating adaptation occurred. Subjects also took longer steps at left heel strike and took narrower steps and shifted their XCOM backward at left toe off (early to late perturbed: BOS_ap, lhs_ increased, p = 0.01, Cohen’s d = 1.09; BOS_ml, lto_ decreased, p = 0.001, Cohen’s d = 1.37; XCOM_ap, lto_ decreased, p < 0.001, Cohen’s d = 1.77) as they adapted. MOS_ap, lhs_ continued trending to pre level during the post block (early post to late post, p = 0.02, Cohen’s d = 1.02).

**Fig. 4.**
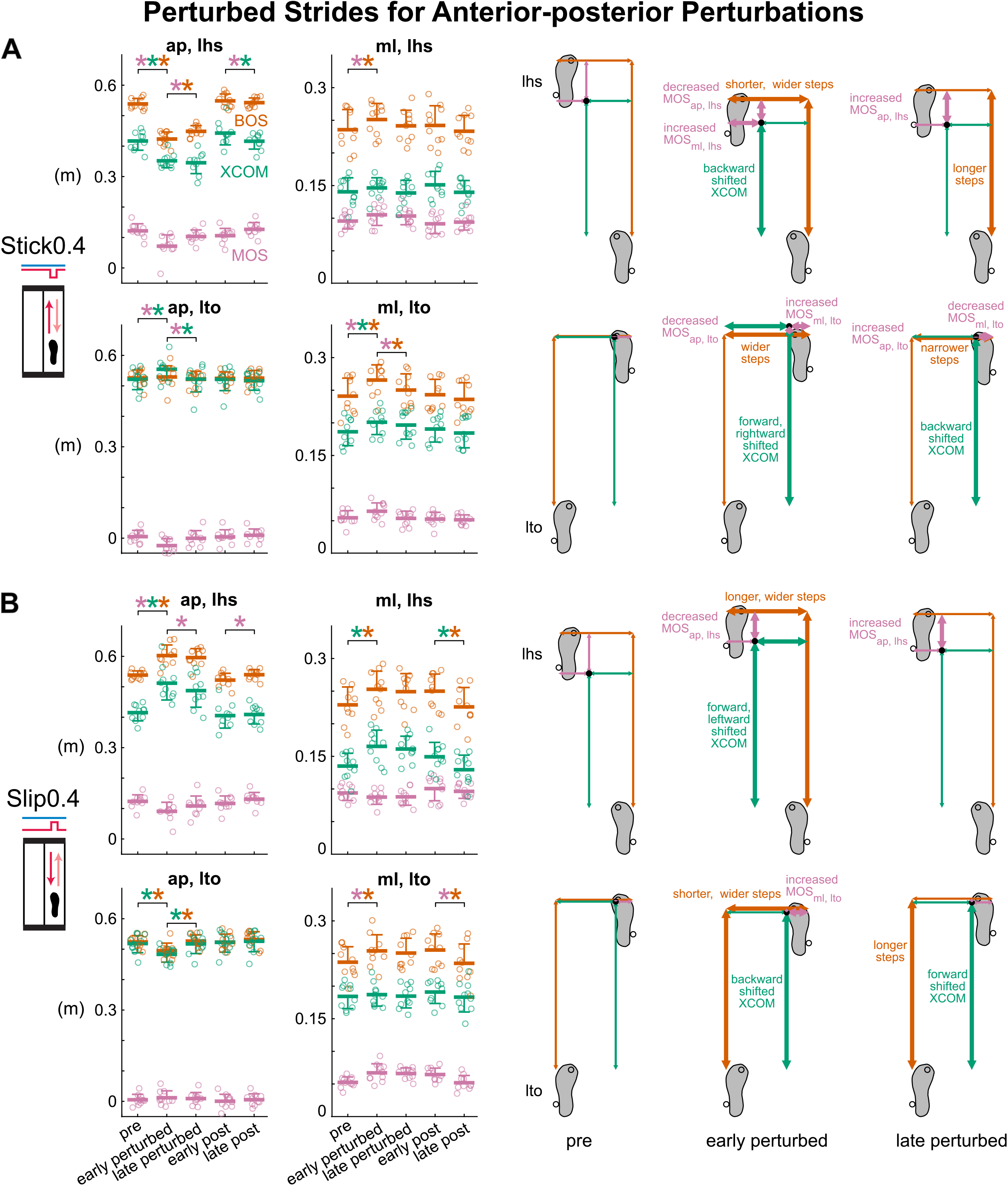
Group-averaged (mean and standard deviation indicated by thick horizontal lines and one-sided error bar, n = 10) anterior-posterior and medial-lateral MOS (pink), BOS (orange), and XCOM (green) at left heel strike and left toe off for pre, early perturbed, late perturbed, early post, and late post during the anterior-posterior perturbations and the corresponding footprints with metrics at pre, early perturbed, and late perturbed. Thick arrows indicate significant changes from the previous phase. In the group-averaged plots, circles are individual subjects. **(A)** Subjects were destabilized at early perturbed then adapted by late perturbed during the Stick0.4 perturbation. **(B)** Subjects were destabilized and adapted during the Slip0.4 perturbation. *: p < 0.05 and color-coded for the specific metrics with the significant differences between phases indicated by the brackets.

Like the Stick0.4 perturbation, subjects adapted to the Slip0.4 perturbation at both left heel strike and left toe off after initially being disrupted (Fig. 4B and Supplemental Fig. S1B). Phase had a significant effect on all metrics except for MOS_ap, lto_ and XCOM_ml, lto_ for the Slip0.4 perturbation (see Supplemental Table S1 for metric specific F, p, and η_p_^2^). Upon initially experiencing the perturbations, anterior-posterior gait stability was disrupted as MOS_ap, lhs_ decreased significantly by 26.6% (p < 0.001, Cohen’s d = 2.34). Subjects also immediately took longer and wider steps and shifted their XCOM forward and leftward at left heel strike (pre to early perturbed: BOS_ap, lhs_ increased, p < 0.001, Cohen’s d = 2.92; BOS_ml, lhs_ increased, p = 0.02, Cohen’s d = 1.05; XCOM_ap, lhs_ increased, p < 0.001, Cohen’s d = 3.39; XCOM_ml, lhs_ increased, p < 0.001, Cohen’s d = 1.53). At left toe off, subjects also immediately increased their step width and shifted their XCOM backward in anticipation of the impending perturbation at mid-stance (pre to early perturbed: BOS_ml, lto_ increased, p = 0.02, Cohen’s d = 1.02; XCOM_ap, lto_ decreased, p < 0.001, Cohen’s d = 1.98). The MOS_ap, lhs_ increased significantly by 19.6% (p = 0.003, Cohen’s d = 1.27) and trended toward pre level from early to late perturbed during adaptation. Subjects also took longer steps and shifted their XCOM forward at left toe off (early to late perturbed: BOS_ap, lto_ increased, p < 0.001, Cohen’s d = 1.49; XCOM_ap, lto_ increased, p < 0.001, Cohen’s d = 1.92). From early post to late post, MOS_ap, lhs_ continued trending to pre level (p = 0.02, Cohen’s d = 1.01).

### 3.3. Adaptation to small medial-lateral perturbations applied on a stride-by-stride basis (hypothesis 1)

The L1 perturbation was initially disruptive, but subjects adapted with more experience with the perturbation (Fig. 5A and Supplemental Fig. S1C). Phase had a significant effect on all metrics except for XCOM_ap, lto_ and MOS_ml, lto_ for the L1 perturbation (see Supplemental Table S1 for metric specific F, p, and η_p_^2^). After initial exposure to the perturbations, MOS_ml, lhs_ was disrupted and increased significantly by 47.3% (p < 0.001, Cohen’s d = 4.27). Subjects also immediately took shorter and wider steps and shifted their XCOM backward and rightward at left heel strike (pre to early perturbed: BOS_ap, lhs_ decreased, p < 0.001, Cohen’s d = 1.46; BOS_ml, lhs_ increased, p = 0.008, Cohen’s d = 1.14; XCOM_ap, lhs_ decreased, p = 0.004, Cohen’s d = 1.22; XCOM_ml, lhs_ decreased, p < 0.001, Cohen’s d = 1.84). Starting with the second L1 perturbation, subjects showed anticipatory responses based on their wider steps at left toe off (pre to early perturbed: BOS_ml, lto_ increased, p = 0.005, Cohen’s d = 1.20). From early to late perturbed, MOS_ml, lhs_ decreased significantly by 8.0% (p = 0.02, Cohen’s d = 1.06) and trended toward pre level, indicating adaptation to the L1 perturbation. Subjects also took longer and narrower steps at both left heel strike and left toe off (early to late perturbed: BOS_ap, lhs_ increased, p = 0.01, Cohen’s d = 1.09; BOS_ml, lhs_ decreased, p = 0.05, Cohen’s d = 0.91; BOS_ap, lto_ increased, p = 0.02, Cohen’s d = 1.02; BOS_ml, lto_ decreased, p = 0.004, Cohen’s d = 1.22) as they adapted. From early post to late post, MOS_ml, lhs_ did not change significantly (p = 0.87, Cohen’s d = 0.30), suggesting that subjects rapidly restored pre level of MOS_lhs_ preference in the medial-lateral direction during the first few strides of the post block.

**Fig. 5.**
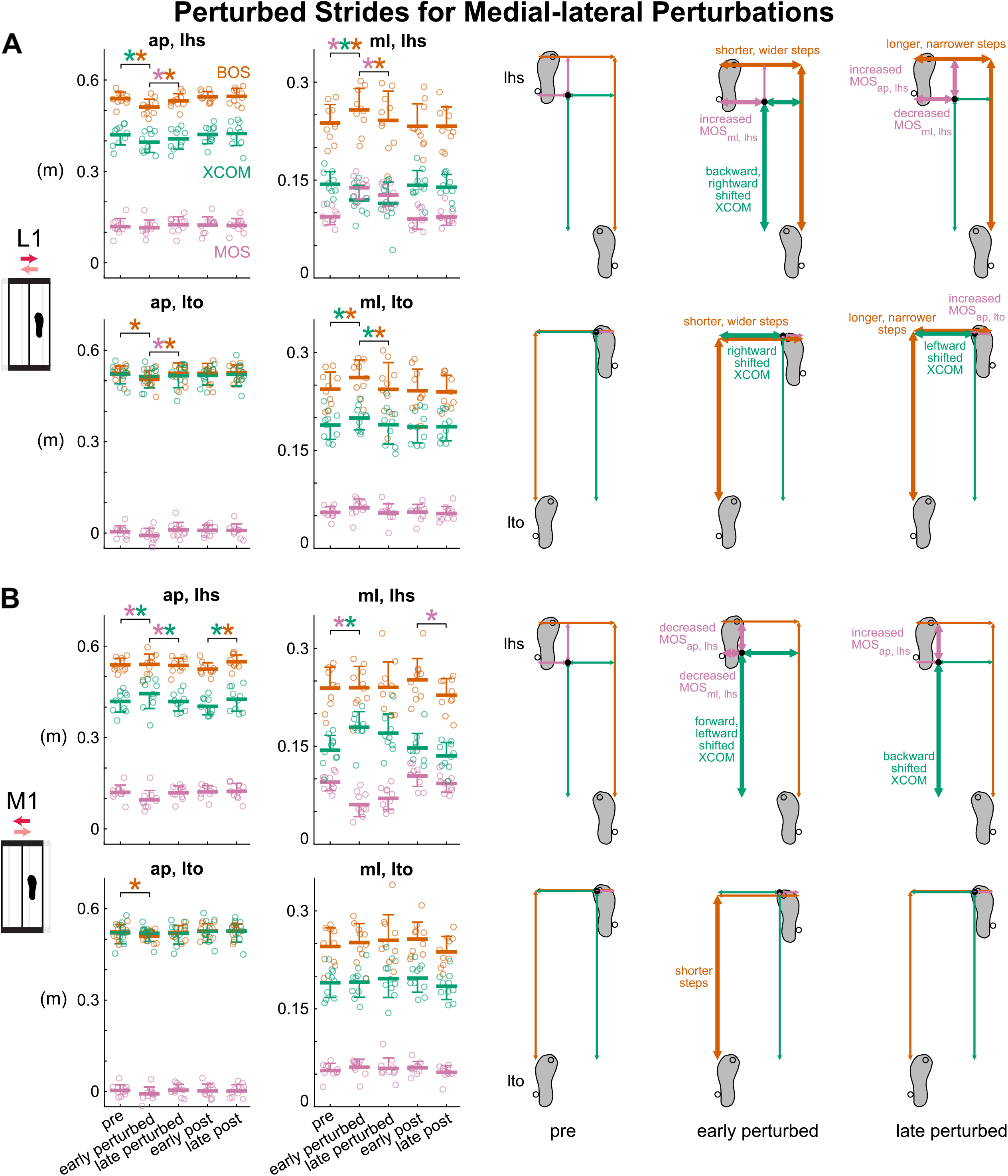
Group-averaged (mean and standard deviation indicated by thick horizontal lines and one-sided error bar, n = 10) anterior-posterior and medial-lateral MOS (pink), BOS (orange), and XCOM (green) at left heel strike and left toe off for pre, early perturbed, late perturbed, early post, and late post during the medial-lateral perturbations and the corresponding footprints with metrics at pre, early perturbed, and late perturbed. Thick arrows indicate significant changes from the previous phase. In the group-averaged plots, circles are individual subjects. **(A)** Subjects adapted to the L1 perturbation after initially being disrupted. **(B)** Subjects adapted to the M1 perturbation after initially being disrupted. *: p < 0.05 and color-coded for the specific metrics with the significant differences between phases indicated by the brackets.

Subjects also adapted to the M1 perturbation that initially disrupted gait stability (Fig. 5B and Supplemental Fig. S1D). Phase had a significant effect on MOS_ap, lhs_, BOS_ap, lhs_, XCOM_ap, lhs_, MOS_ml, lhs_, XCOM_ml, lhs_, and BOS_ap, lto_ for the M1 perturbation (see Supplemental Table S1 for metric specific F, p, and η_p_^2^). Upon initially responding to the perturbations, gait stability was disrupted in both directions as MOS_ap, lhs_ decreased significantly by 20.3% (p = 0.002, Cohen’s d = 1.28) and MOS_ml, lhs_ decreased significantly by 36.3% (p < 0.001, Cohen’s d = 2.80). After the initial perturbation, subjects immediately shifted their XCOM forward and leftward at left heel strike (pre to early perturbed: XCOM_ap, lhs_ increased, p = 0.01, Cohen’s d = 1.09; XCOM_ml, lhs_ increased, p < 0.001, Cohen’s d = 1.88). At left toe off, subjects also immediately took shorter steps (pre to early perturbed: BOS_ap, lto_ decreased, p = 0.04, Cohen’s d = 0.94). Subjects adapted after the initial disruption as MOS_ap, lhs_ increased significantly by 23.9% (p = 0.004, Cohen’s d = 1.21) and trended toward pre level by late perturbed. Subjects also shifted their XCOM backward at left heel strike (early to late perturbed: XCOM_ap, lhs_ decreased, p = 0.01, Cohen’s d = 1.11).

### 3.4. Which component of margin of stability was affected more by the small perturbations?

During disruption, perturbations had different effects on the magnitudes of the changes in the BOS and XCOM at left heel strike (Fig. 6A) and left toe off (Supplemental Fig. S2A). For the Stick0.4 perturbation, the BOS_lhs_ was disrupted more than XCOM_lhs_ based on the larger changes in the BOS_lhs_ compared to the changes in XCOM_lhs_ that were significantly different in the anterior-posterior direction and trending towards significance in the medial-lateral direction (|ΔBOS_ap, lhs_| > |ΔXCOM_ap, lhs_| p = 0.001, Cohen’s d = 1.60); |ΔBOS_ml, lhs_| > |ΔXCOM_ml, lhs_| p = 0.08, Cohen’s d = 0.62). For the Slip0.4 and M1 perturbations, the XCOM_lhs_ was disrupted more than BOS_lhs_ as the change in XCOM_lhs_ was greater than the change in BOS_lhs_ in both directions (Slip0.4: |ΔXCOM_ap, lhs_| > |ΔBOS_ap, lhs_| p < 0.001, Cohen’s d = 2.00; |ΔXCOM_ml, lhs_| > |ΔBOS_ml, lhs_| p = 0.04, Cohen’s d = 0.73; M1: |ΔXCOM_ap, lhs_| > |ΔBOS_ap,_ lhs| p = 0.06, Cohen’s d = 0.68; |ΔXCOM_ml, lhs_| > |ΔBOS_ml, lhs_| p < 0.001, Cohen’s d = 1.93). For the L1 perturbation, there was no difference between the changes in BOS_lhs_ and XCOM_lhs_ in either direction (p’s > 0.21, Cohen’s d’s < 0.43).

**Fig. 6.**
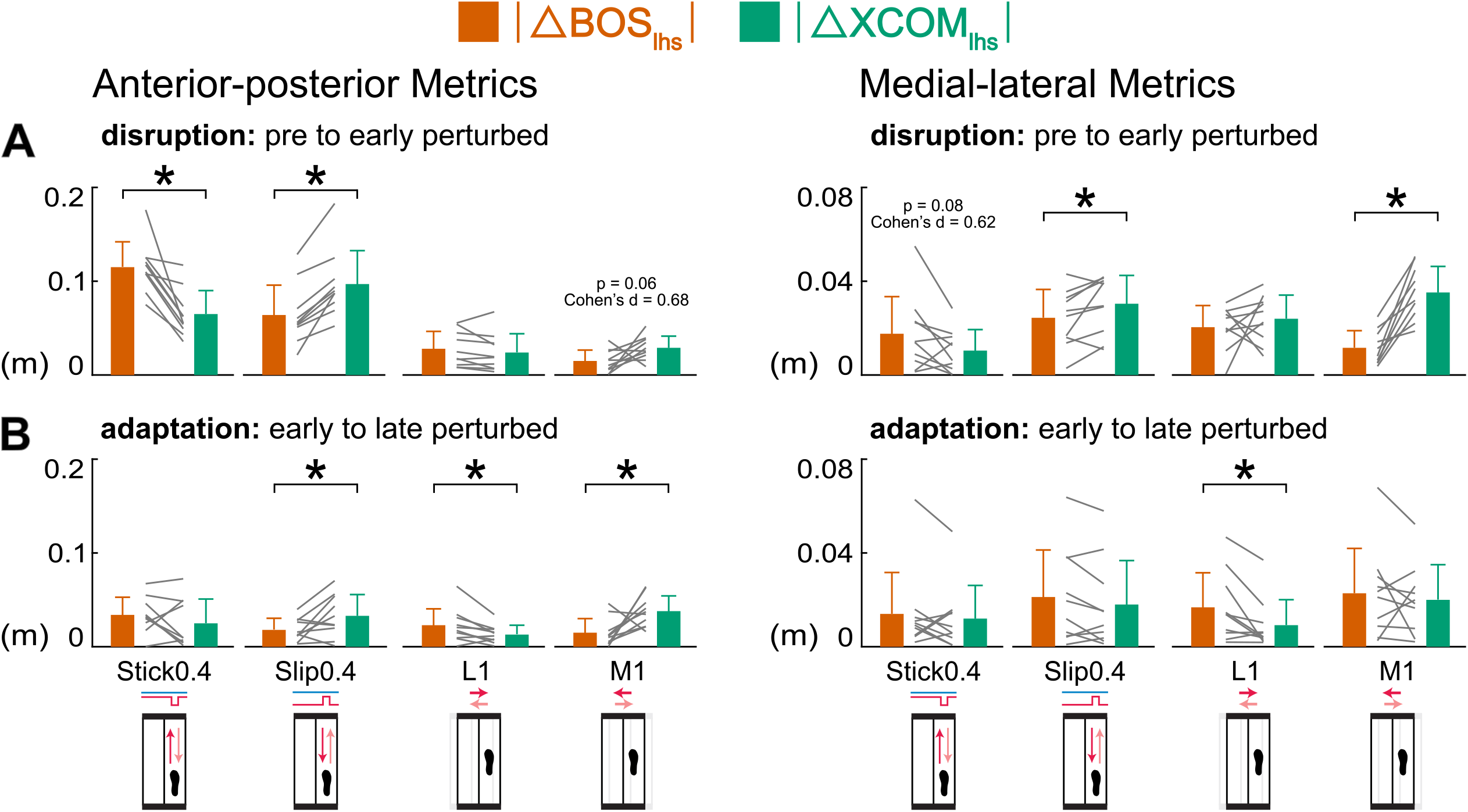
Group-averaged (mean and standard deviation, n = 10) |ΔBOS_lhs_| (orange) compared with |ΔXCOM_lhs_| (green) during disruption (pre to early perturbed) and adaptation (early to late perturbed). Gray lines between bars are individual subjects. **(A)** During disruption, BOS_lhs_ was disrupted more by the Stick0.4 perturbation, XCOM_lhs_ was disrupted more by the Slip0.4 and M1 perturbations. **(B)** During adaptation, BOS_ap, lhs_ adapted more in the L1 perturbation, XCOM_ap, lhs_ adapted more in the Slip0.4 and M1 perturbations, BOS_ml, lhs_ adapted more in the L1 perturbation. *: p < 0.05 and significant differences between |ΔBOS_lhs_| and |ΔXCOM_lhs_|.

### 3.5. Which component contributed more to the adaptation of margin of stability?

During adaptation, the component that was the primary contributor to the adaptation of MOS at left heel strike (Fig. 6B) and left toe off (Supplemental Fig. S2B) depended on the perturbation. The |ΔBOS_ap, lhs_| was larger than the |ΔXCOM_ap, lhs_| for the L1 perturbation (p = 0.03, Cohen’s d = 0.85). The |ΔXCOM_ap, lhs_| was greater than the |ΔBOS_ap,_ lhs| for the Slip0.4 and M1 perturbations (p’s < 0.05, Cohen’s d’s > 0.72), suggesting that XCOM_ap, lhs_ adapted more than BOS_ap, lhs_ during those perturbations. There was no significant difference between |ΔBOS_ap, lhs_| and |ΔXCOM_ap,_ lhs| in the Stick0.4 perturbation (p = 0.27, Cohen’s d = 0.37). For the medial-lateral metrics, the only significant difference was the larger |ΔBOS_ml, lhs_| compared to the |ΔXCOM_ml, lhs_| for the L1 perturbation (p = 0.02, Cohen’s d = 0.90), suggesting that BOS_ml, lhs_ adapted more in the lateral shift perturbation. There were no significant differences between the |ΔBOS_ml, lhs_| and |ΔXCOM_ml, lhs_| in the other perturbations (p’s > 0.12, Cohen’s d’s < 0.53).

### 3.6. Did subjects use more feedforward strategies when adapting to small perturbations applied on a stride-by-stride basis? (hypothesis 2)

Catch strides revealed subjects used feedforward strategies upon initially experiencing the anterior-posterior perturbations but did not use more anticipatory control as adaptation progressed (Fig. 7A, B and Supplemental Fig. S1A, B). For the Stick0.4 perturbation, phase had a significant effect on all metrics except for BOS_ap, lto_, BOS_ap, lhs_, MOS_ml, lhs_ while for the Slip0.4 perturbation phase had a significant effect on MOS_ml, lto_, BOS_ml, lto_, MOS_ap, lhs_, XCOM_ap, lhs_, MOS_ml, lhs_, BOS_ml, lhs_, and XCOM_ml, lhs_ (see Supplemental Table S4 for metric specific F, p, and η_p_^2^). At early catch, MOS_ap, lto_ and MOS_ml, lto_ deviated significantly from their pre levels for the Stick0.4 (MOS_ap, lto_ decreased, p = 0.001, Cohen’s d = 1.44; MOS_ml, lto_ increased, p = 0.02, Cohen’s d = 0.96) and the Slip0.4 (MOS_ml, lto_ increased, p = 0.004, Cohen’s d = 1.20) perturbations. Subjects also took wider steps for both perturbations (pre to early catch: BOS_ml, lto_ increased, p’s < 0.003, Cohen’s d’s > 1.25) and shifted their XCOM forward and rightward at left toe off for the Stick0.4 perturbation (pre to early catch: XCOM_ap, lto_ increased, p = 0.02, Cohen’s d = 0.99; XCOM_ml, lto_ increased, p = 0.009, Cohen’s d = 1.07). At left heel strike for the Stick0.4 perturbation, subjects marginally shifted their XCOM forward to 0.44 ± 0.05 m (pre to early catch: XCOM_ap, lhs_, p = 0.130, Cohen’s d = 0.65), which was in the opposite direction of the perturbed response of 0.35 ± 0.02 m, revealing evidence of anticipating an impending perturbation. MOS_ml. lto_ and MOS_ap, lhs_ trended to their pre levels by late catch for the Stick0.4 perturbation (early to late catch: MOS_ml. lto_ decreased, p = 0.02, Cohen’s d = 0.96; MOS_ap, lhs_ increased, p = 0.02, Cohen’s d = 0.99). Subjects also took narrower steps for the Stick0.4 perturbation (early to late catch: BOS_ml,_ lto decreased, p = 0.003, Cohen’s d = 1.22; BOS_ml, lhs_ decreased, p = 0.007, Cohen’s d = 1.10) and shifted their XCOM rightward at left heel strike for both perturbations (early to late catch: XCOM_ml, lhs_ decreased, p’s < 0.007, Cohen’s d’s > 1.11). Although MOS_ml, lto_ and MOS_ml, lhs_ increased significantly from pre to late catch for the Slip0.4 perturbation (MOS_ml, lto_, p = 0.03, Cohen’s d = 0.91; MOS_ml, lhs_, p = 0.03, Cohen’s d = 0.88), there were no significant differences between early catch and late catch (MOS_ml, lto_, p = 0.64, Cohen’s d = 0.29; MOS_ml, lhs_, p = 0.11, Cohen’s d = 0.67), suggesting that subjects did not use more feedforward strategies as they experienced more anterior-posterior perturbations.

**Fig. 7.**
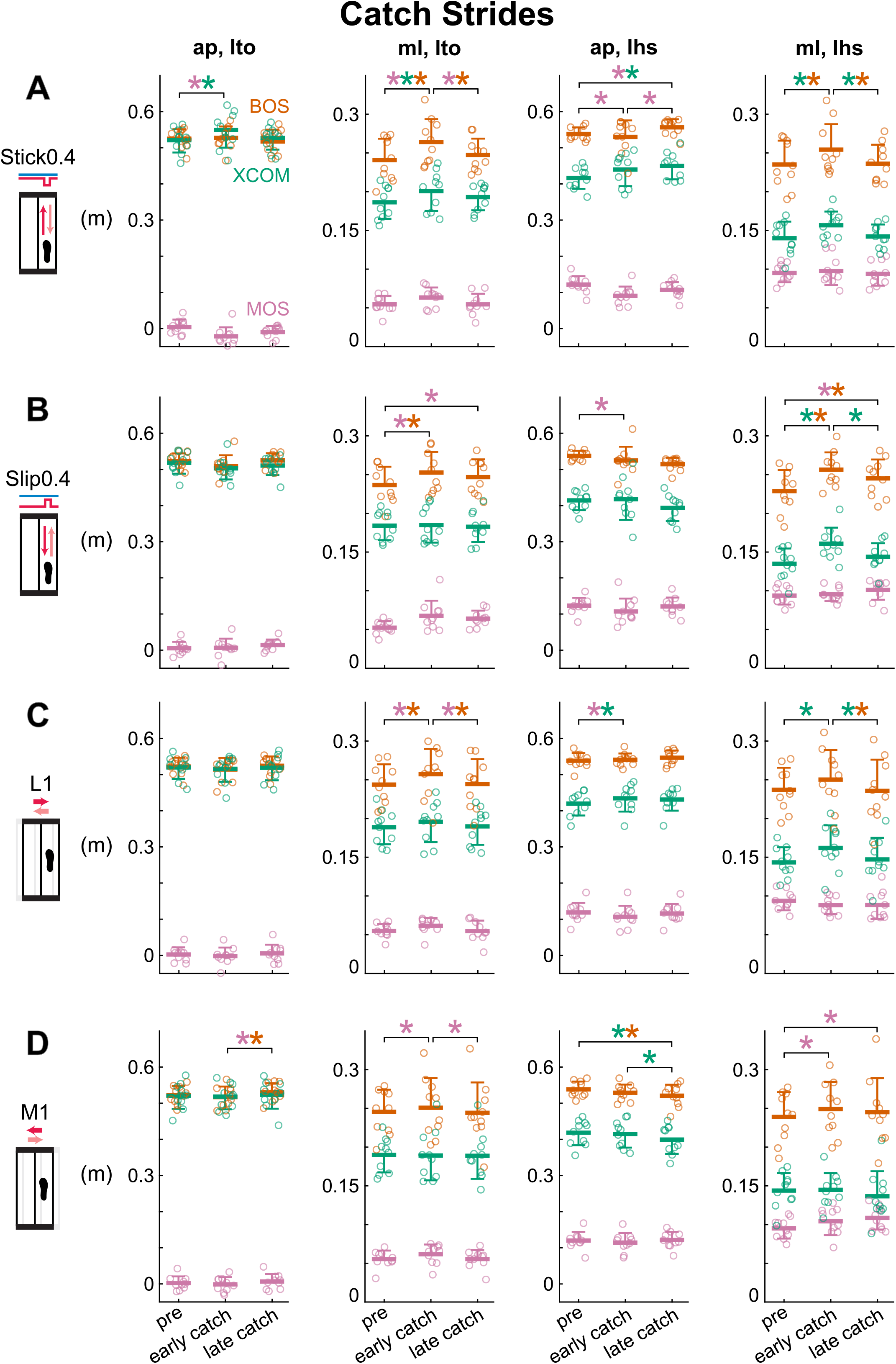
Group-averaged (mean and standard deviation indicated by thick horizontal lines and one-sided error bar, n = 10) anterior-posterior and medial-lateral MOS (pink), BOS (orange), and XCOM (green) at left heel strike and left toe off for pre, early catch, late catch for the Stick0.4, Slip0.4, L1, and M1 perturbations. Circles are individual subjects. **(A)** Stick0.4 perturbation, **(B)** Stick0.4 perturbation, **(C)** L1 perturbation **(D)** M1 perturbation induced anticipatory responses at early catch, but subjects did not use more feedforward strategies by late catch. *: p < 0.05 and color-coded for the specific metrics with the significant differences between phases indicated by the brackets.

Based on the responses to the catches, subjects employed feedforward strategies after the initial medial-lateral perturbations, however, more feedforward strategies were not used during adaptation (Fig. 7C, D and Supplemental Fig. S1C, D). Phase had a significant effect on MOS_ml, lto_, BOS_ml, lto_, MOS_ap, lhs_, XCOM_ap, lhs_, BOS_ml, lhs_, and XCOM_ml,_ lhs for the L1 perturbation and MOS_ap, lto_, BOS_ap, lto_, MOS_ml, lto_, BOS_ap, lhs_, XCOM_ap, lhs_, and MOS_ml, lhs_ for the M1 perturbation (see Supplemental Table S4 for metric specific F, p, and η_p_^2^). From pre to early catch, MOS_ml, lto_ increased significantly for both perturbations (p’s < 0.02, Cohen’s d > 1.04). Subjects also took wider steps at left toe off for the L1 perturbation (pre to early catch: BOS_ml, lto_ increased, p = 0.004, Cohen’s d = 1.20). At left heel strike, subjects shifted their XCOM forward and leftward for the L1 perturbation (pre to early catch: XCOM_ap, lhs_ increased, p = 0.04, Cohen’s d = 0.85; XCOM_ml, lhs_ increased, p = 0.002, Cohen’s d = 1.28) and increased their MOS_ml, lhs_ for the M1 perturbation (pre to early catch, p = 0.002, Cohen’s d = 1.27), these trends were in the opposite direction of the responses for pre to early perturbed. These results suggest that subjects had anticipatory behavior after the initial medial-lateral perturbations. From early to late catch, MOS_ml. lto_ decreased significantly for both perturbations (p’s < 0.009, Cohen’s d’s > 1.08). Subjects also took narrower steps at left toe off and shifted their XCOM rightward at left heel strike for the L1 perturbation (early to late catch: BOS_ml, lto_ decreased, p = 0.006, Cohen’s d = 1.13; XCOM_ml, lhs_ decreased, p = 0.01, Cohen’s d = 1.02). For the M1 perturbation, MOS_ml, lhs_ increased and subjects took shorter steps and shifted their XCOM rightward at left heel strike (pre to late catch: MOS_ml, lhs_ increased, p < 0.0001, Cohen’s d = 1.88; BOS_ap, lhs_ decreased, p = 0.01, Cohen’s d = 1.06; XCOM_ap, lhs_ decreased, p = 0.004, Cohen’s d = 1.20), whereas no significant differences were found between their early catch and late catch except for XCOM_ap, lhs_ (early to late catch: MOS_ml, lhs_, p = 0.16, Cohen’s d = 0.61; BOS_ap, lhs_, p = 0.29, Cohen’s d = 0.49; XCOM_ap, lhs_ decreased, p = 0.02, Cohen’s d = 0.96). These results suggest that subjects generally did not use more feedforward strategies as they adapted to the medial-lateral perturbations.

### 3.7. Are margin of stability and its components sensitive to directions of small perturbations? (hypothesis 3)

Anterior-posterior metrics at left heel strike were more sensitive to the anterior-posterior perturbations when subjects were disrupted at early perturbed but not during adaptation from early to late perturbed (Fig. 8A). During disruption, perturbation direction had a significant effect on each anterior-posterior metric at left heel strike (F’s (3,27) > 7.92, p’s < 0.002, η_p_^2^’s > 0.46). All perturbations produced a decrease in MOS_ap, lhs_, during disruption of which, the decreases for the Stick0.4 and Slip0.4 perturbations were observed in all subjects and significantly larger than the decrease for the L1 perturbation (p’s < 0.04, Cohen’s d’s > 0.94). For early perturbed, the BOS_ap, lhs_ and XCOM_ap, lhs_ had the largest decrease with the Stick0.4 perturbation compared to the decrease with the L1 perturbation (p’s < 0.02, Cohen’s d’s > 1.03). The BOS_ap, lhs_ and XCOM_ap, lhs_ had the largest increase with the Slip0.4 perturbation compared to the increase with the M1 perturbation for early perturbed (p’s < 0.001, Cohen’s d’s > 1.72). During adaptation, all perturbations resulted in an increased MOS_ap, lhs_ of similar magnitude (F (3,27) = 1.36, p = 0.28, η_p_^2^ = 0.13). The Stick0.4 and L1 perturbations resulted in increases of BOS_ap, lhs_ during adaptation, whereas the only significant difference was between Stick0.4 and Slip0.4 perturbations mainly due to differences in sign (p = 0.04, Cohen’s d = 0.92). Conversely, although perturbation direction did not have a significant main effect on XCOM_ap, lhs_ (F (3,27) = 2.83, p = 0.06, η_p_^2^ = 0.24), the Slip0.4 and M1 perturbations resulted in decreases of XCOM_ap, lhs_ during adaptation.

**Fig. 8.**
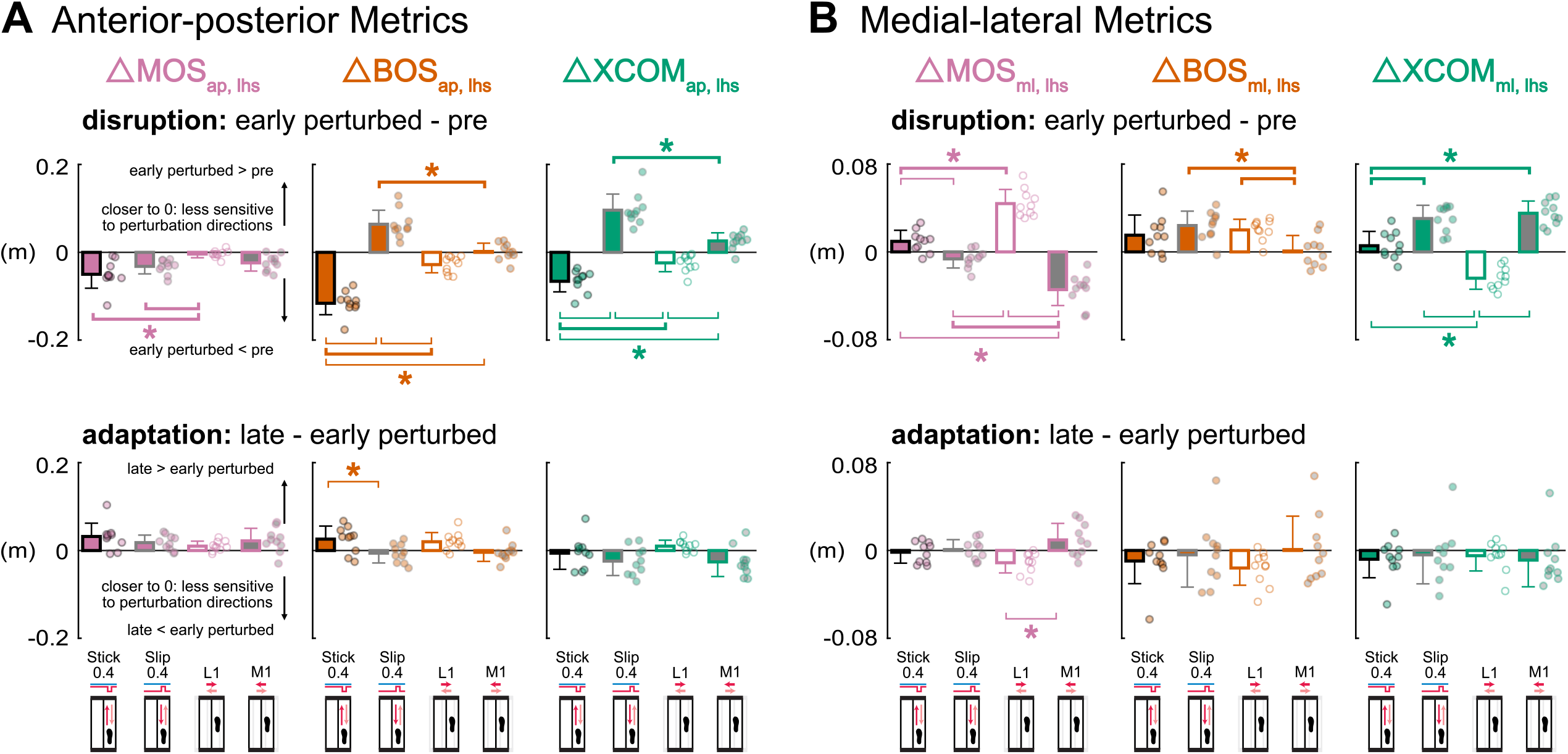
Group-averaged (mean and standard deviation, n = 10) ΔMOS_lhs_ (pink), ΔBOS_lhs_ (orange), ΔXCOM_lhs_ (green) during disruption (early perturbed minus pre) and adaptation (late minus early perturbed) for the Stick0.4 (black outline), Slip0.4 (gray outline), L1 (white fill), and M1 (gray fill) perturbations. Circles are individual subjects. **(A)** Anterior-posterior metrics at left heel strike were more sensitive to the anterior-posterior perturbations during disruption but not adaptation. **(B)** MOS_ml, lhs_ and XCOM_ml, lhs_ were more sensitive to the medial-lateral perturbations during disruption but not adaptation. * with thick brackets: p < 0.05 and significant same signed differences between two perturbation conditions, * with thin brackets: p < 0.05 and significant opposite signed differences between two perturbation conditions.

The medial-lateral metrics at left heel strike, MOS_ml, lhs_ and XCOM_ml, lhs_, were more sensitive to the medial-lateral perturbations during disruption but not during adaptation (Fig. 8B). During disruption, perturbation direction had a significant effect on each medial-lateral metric at left heel strike (F’s (3,27) > 5.32, p’s < 0.006, η_p_^2^’s > 0.36). For early perturbed, all subjects increased and decreased MOS_ml, lhs_ in the L1 and M1 perturbations, respectively. MOS_ml, lhs_ had the largest increase with the L1 perturbation compared to the increase with the Stick0.4 perturbation (p < 0.001, Cohen’s d = 2.07), while the MOS_ml, lhs_ had the largest decrease with the M1 perturbation compared to the decrease with the Slip0.4 perturbation (p < 0.001, Cohen’s d = 1.67). All perturbations produced an increase in BOS_ml, lhs_, during disruption of which, the increase for the M1 perturbation was significantly smaller than the increases for the Slip0.4 and L1 perturbations (p’s < 0.03, Cohen’s d’s > 0.98). For the XCOM_ml, lhs_ for early perturbed, the increase of XCOM_ml, lhs_ for all subjects in the M1 perturbation was significantly larger than the increase in the Stick0.4 perturbation (p < 0.001, Cohen’s d = 1.79), while the decrease of XCOM_ml, lhs_ for all subjects in the L1 perturbation was significantly different than the increases in the Stick0.4 and Slip0.4 perturbations (p’s < 0.001, Cohen’s d’s > 1.83). During adaptation, perturbation direction had a significant effect on ΔMOS_ml, lhs_ (F (3,27) = 5.89, p = 0.003, η_p_^2^ = 0.40). The increase of MOS_ml, lhs_ in the M1 perturbation was significantly different than the decrease in the L1 perturbation (p = 0.002, Cohen’s d = 1.32), largely due to differences in sign. For BOS_ml,_ lhs and XCOM_ml, lhs_, nearly all perturbations resulted in a decrease of similar magnitudes during adaptation (F’s (3,27) < 0.93, p’s > 0.44, η_p_^2^’s < 0.10).

### 3.8. Do responses in margin of stability and its components scale with perturbation size? (hypothesis 4)

Anterior-posterior metrics at left heel strike scaled with perturbation size when subjects were disrupted at early perturbed but not during adaptation from early to late perturbed (Fig. 9A). Perturbation size had a main effect on the changes during disruption for each anterior-posterior metric at left heel strike (F’s (1,27) > 9.51, p’s < 0.006, η_p_^2^’s > 0.25). There was a significant interaction effect between perturbation size and perturbation condition for the changes in BOS_ap, lhs_ during disruption (F (1,27) = 5.53, p = 0.03, η_p_^2^ = 0.17). Paired t-tests revealed that the half-size perturbations produced significantly smaller magnitude changes in BOS_ap, lhs_ than the regular perturbations within each perturbation condition (Stick0.2 vs. Stick0.4 p = 0.001, Cohen’s d = 1.61; Slip0.2 vs. Slip0.4 p = 0.01, Cohen’s d = 1.01). These results suggest that subjects scaled BOS_ap, lhs_ more in the stick perturbations than slip perturbations at early perturbed. During adaptation, perturbation size had a significant effect on |ΔBOS_ap, lhs_| (F (1,27) = 4.75, p = 0.04, η_p_^2^ = 0.15) but not on |ΔMOS_ap, lhs_ | and |ΔXCOM_ap, lhs_ | (F’s (1,27) < 1.70, p’s > 0.20, η_p_^2^’s < 0.07), indicating most anterior-posterior metrics at left heel strike did not scale with perturbation size during adaptation.

**Fig. 9.**
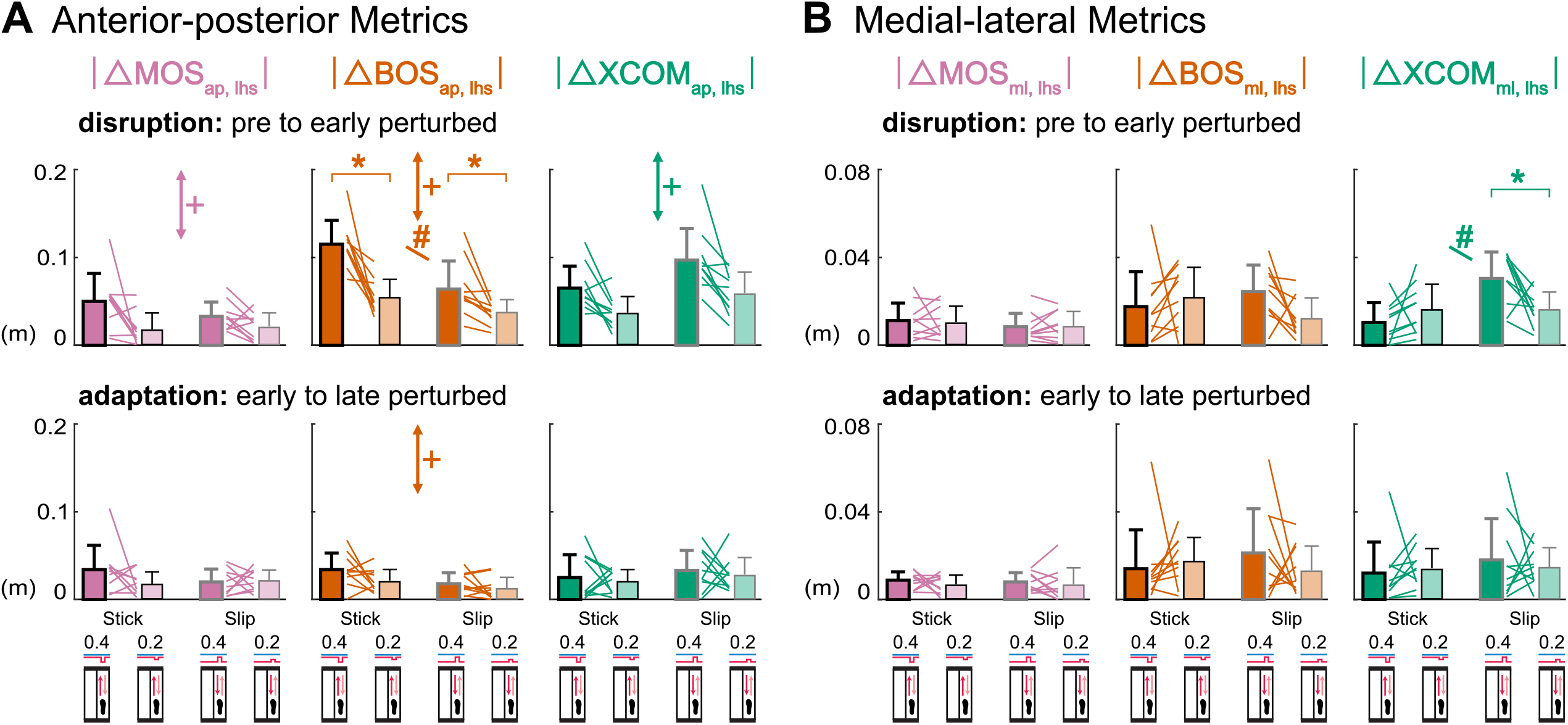
Group-averaged (mean and standard deviation, n = 10) |ΔMOS_lhs_| (pink), |ΔBOS_lhs_| (orange), |ΔXCOM_lhs_| (green) during disruption (pre to early perturbed) and adaptation (early to late perturbed) for the anterior-posterior perturbations. Dark bars indicate the regular size perturbations (Stick0.4 and Slip 0.4) and light bars indicate the half-size perturbations (Stick0.2 and Slip 0.2). Colored lines between bars are individual subjects. **(A)** Anterior-posterior metrics at left heel strike scaled with perturbation size during disruption but not adaptation. **(B)** Medial-lateral metrics at left heel strike did not scale with perturbation size during disruption and adaptation. +: p < 0.05, perturbation size effect (perturbation condition effect was not reported). #: p < 0.05, interaction effect between perturbation size (regular, 0.4; half-size, 0.2) and perturbation condition (stick, slip). *: p < 0.025, significant differences between the regular and half-size perturbations of same condition.

Medial-lateral metrics at left heel strike did not scale with perturbation size when subjects were disrupted or during adaptation (Fig. 9B). Perturbation size did not have a main effect on the changes during disruption for any medial-lateral metric at left heel strike (F’s (1,27) < 1.19, p’s > 0.28, η_p_^2^’s < 0.05). Despite the significant interaction effect on |ΔXCOM_ml, lhs_| during disruption (F (1,27) = 7.96, p = 0.009, η_p_^2^ = 0.23), paired t-tests only detected significant differences between Slip perturbations (p = 0.02, Cohen’s d = 0.93) but not between Stick perturbations (p = 0.21, Cohen’s d = 0.43). For the changes of each medial-lateral metric at left heel strike during adaptation, there was neither a main effect of perturbation size nor a significant interaction effect (F’s (1,27) < 1.23, p’s > 0.27, η_p_^2^’s < 0.05). These results suggest that medial-lateral metrics at left heel strike did not scale with perturbation size during disruption or adaptation.

## 4. Discussion

We sought to determine how small magnitude multidirectional perturbations affect modulation of margin of stability, base of support, and extrapolated center of mass on a stride-by-stride basis. In the anterior-posterior direction, subjects mainly modulated XCOM_ap, lhs_ when adapting to the belt acceleration (Slip0.4) and medial shift (M1) perturbations. In the medial-lateral direction, subjects primarily modulated BOS_ml, lhs_ when adapting to the lateral shift (L1) perturbation. Metrics at left toe off and catch strides revealed that subjects immediately took wider steps for the Stick0.4, Slip0.4, and L1 perturbations and shifted their XCOM_ap, lto_ forward for the Stick0.4 perturbation before perturbation onset, providing evidence of feedforward strategies. Despite the combination of feedback and feedforward strategies upon initial exposure to the perturbations, subjects did not use more feedforward strategies during adaptation. When first experiencing the perturbations, anterior-posterior metrics at left heel strike for perturbed strides were more sensitive to perturbations in the coincident anterior-posterior direction, and likewise, MOS_ml, lhs_ and XCOM_ml, lhs_ were more sensitive to perturbations in the coincident medial-lateral direction. Additionally, anterior-posterior metrics at left heel strike scaled with perturbation size upon initially responding to the perturbations (at disruption). These findings suggest that applying small perturbations on a stride-by-stride basis led to the adaptation of the margin of stability, providing new insights regarding the extent of locomotor adaptation to small perturbations. Further, perturbation directions could be used to target specific aspects of gait stability control (e.g., primarily modulating base of support or center of mass).

One of the main findings was that small magnitude perturbations altered gait stability, i.e., the margin of stability at left heel strike, in specific directions. For most perturbations, the MOS_ap, lhs_ and MOS_ml, lhs_ initially decreased before adapting back to pre levels (Fig. 4, 5), suggesting that subjects were destabilized initially before gradually regaining stability during the Stick0.4, Slip0.4, and M1 perturbations. These results are consistent with previous studies that showed that treadmill belt and shift perturbations decreased MOS_ap_ (11, 25, 42). Adjusting foot placement and BOS is a common approach to modulate MOS (6, 23), which was evident in our Stick0.4 perturbation. Other studies demonstrated the importance of center of mass control to modulate MOS (11, 30), which was evident in the Slip0.4 and M1 perturbations. MOS_lhs_ did not always decrease in response to perturbations. For the Stick0.4 and L1 perturbation, the MOS_ml, lhs_ initially increased, however, instead of decreasing as seen in the other perturbations. This increased MOS_ml, lhs_ likely resulted from a combination of a more lateral (leftward) foot placement (i.e., increased BOS_ml, lhs_) and redirected XCOM_lhs_ rightward during the lateral shift perturbation. In another study that applied lateral shift perturbations at early single leg support, people used the gluteus medius of their swing leg to place their subsequent foot more outward (9) in the direction of the potential fall (27, 43), which would help increase the MOS_ml_ to maintain a more stable body position (19). Larger MOS_ml_ often occurs in response to medial-lateral treadmill shift perturbations (42, 44, 45), similar to the increased MOS_ml, lhs_ in our L1 perturbation. Interestingly, subjects maintained this increased MOS_ml, lhs_ for the Stick0.4 perturbation but gradually reduced the MOS_ml, lhs_ for the L1 perturbation, suggesting that subjects either could not or did not have a reason to decrease MOS_ml, lhs_ while gaining more experience with the Stick0.4 perturbation.

At left heel strike, subjects primarily modulated foot placement compared to the extrapolated center of mass in the belt deceleration (during disruption) and lateral shift (during adaptation) perturbations. Researchers are increasingly finding that BOS control is an active neuromechanical balance strategy that is adjusted based on the state of the center of mass during walking (4, 43, 46, 47). Additionally, the central nervous system is actively involved in this foot placement strategy (48, 49). In our study, both the BOS_ap, lhs_ and XCOM_ap, lhs_ decreased in response to the Stick0.4 perturbation, but subjects primarily adjusted BOS_ap, lhs_ by taking shorter steps during disruption. The reduction and smaller adjustment of XCOM_ap, lhs_ compared to the BOS_ap, lhs_ was likely a consequence of the rotation and acceleration of the upper body backward relative to the lower body created by the stick perturbations. The significant BOS_ap, lhs_ adjustment was likely needed to rapidly interrupt the forward progression of the swing leg to compensate for the interrupted progression of the stance leg caused by the Stick0.4 perturbation (26). These reactions conform to the theory of stepping in the direction of the potential fall and to the maintenance of forward progression on the treadmill (27, 43, 50). During adaptation to the Stick0.4 perturbation, there were similar magnitude changes in the BOS_ap, lhs_ and XCOM_ap, lhs_, suggesting a less dominant stepping strategy during adaptation compared to the dominant foot placement strategy during disruption. For the lateral shift perturbation, subjects appeared to employ combination of a foot placement strategy through swing leg control that increased the BOS_ml, lhs_ (9), while also using an ankle strategy with the stance leg that decreased XCOM_ml, lhs_ by a similar magnitude (51). Changes in the BOS_ml, lhs_ then dominated during adaptation to the L1 perturbation. This aligns with the predominant stepping strategy people often employ, taking shorter and wider steps during medial waist-pull perturbations that produced relative movements of the BOS and XCOM similar to our L1 perturbation (52).

Our results also showed that subjects primarily modulated the extrapolated center of mass at left heel strike in the belt acceleration and medial shift perturbations. Both the BOS_ap, lhs_ and XCOM_ap, lhs_ increased in response to our Slip0.4 perturbation, but subjects mainly modulated their XCOM_ap, lhs_ while adapting to the Slip0.4 perturbation. The XCOM_ap, lhs_ responses were likely related to the decreased hip flexor moment of the right stance leg to reduce the forward rotation and acceleration of the upper body (11). Even though the BOS_ap, lhs_ also changed during adaptation, these changes were smaller than the XCOM_ap, lhs_ and were probably a consequence of the facilitated forward swing phase induced by the Slip0.4 perturbation. For the medial shift perturbation, subjects primarily adjusted XCOM_lhs_, interestingly, in the orthogonal anterior-posterior direction, likely by employing an ankle strategy (27) that incorporated increased tibialis anterior activity of the stance leg (51). Subjects barely adjusted the BOS_ap, lhs_, likely due to the small magnitude of our M1 perturbation. In other medial-lateral perturbations, subjects often take crossover steps to adjust BOS in the anterior-posterior direction. (53, 54).

Our results suggest that subjects can flexibly employ balance strategies in response to small magnitude perturbations applied on a stride-by-stride basis, depending on the perturbation direction. When the central nervous system detects a perceived fall, the foot placement strategy shifts the swing foot to the direction of the potential fall, such that at heel strike the undesired movement induced by perturbations could be reversed or at least mitigated (27, 28). Another strategy is adjusting the extrapolated center of mass to stop the potential fall through the stance leg, which is similar to the ankle strategy in other studies (27, 52, 55). The foot placement strategy has the advantage of a large adjustment range, however, it acts slowly due to the time needed for active integration of sensory information from the proprioceptive, visual, and vestibular systems (27, 28). Conversely, the extrapolated center of mass strategy allows for quick responses during single stance, whereas its modulation range is constrained to the relatively small contact area under the stance foot (27, 55, 56). A recent visual perturbation study shows that foot placement and ankle strategies are interdependent and negatively correlated, suggesting that these two strategies complement each other (57). Both strategies may increase the energetic cost as a consequence of the cost of redirecting foot placement and extrapolated center of mass (58–61), but adaptation over time usually leads to a more economical gait (62, 63). Thus, depending on the perturbation direction, people appear to recruit more readily available and efficient balance strategies for gait stability when experiencing small magnitude perturbations applied on a stride-by-stride basis.

In our study, subjects employed a feedforward strategy that involved direction independent foot placement but direction dependent extrapolated center of mass control after being initially perturbed. Subjects immediately took wider steps at left toe off for the early catch strides after experiencing the Stick0.4, Slip0.4, and L1 perturbations, indicating a direction independent anticipatory strategy (Fig. 7A, B, C). These changes reflect anticipatory responses because left toe off occurs immediately before the expected perturbation at right mid-stance. For pre to early perturbed, like the catches, subjects also took wider steps at left toe off (Fig. 4 and 5A), suggesting a general feedforward strategy of increasing step width. Taking wider steps can increase the dynamic stability in the medial-lateral direction (27, 64) and can help prepare for an upcoming perturbation in any direction. For the Stick0.4 perturbation, XCOM_ap, lto_ shifted forward for early catch (Fig. 7A) and early perturbed (Fig. 4A), which was opposite to the potential fall direction of the Stick0.4 perturbation, revealing direction dependent anticipatory XCOM adjustments. Forward shifts of the COM or XCOM position to prepare for impending backward fall inducing perturbations (65–67) and a reduction of forward COM velocity to prepare for impending forward fall inducing perturbations (68, 69) are common anticipatory balance strategies in response to anterior-posterior perturbations. The lack of anterior-posterior XCOM adjustments for the Slip0.4 perturbation was likely due to the less risk of forward fall compared to backward fall (10, 70).

Although subjects initially anticipated, they did not rely more on feedforward strategies when adapting to small magnitude perturbations applied on a stride-by-stride basis. In response to repeated perturbations, people often adapt using feedback (reactive) and feedforward (proactive) control (71, 72). Feedback control relies on sensory detection of unexpected disturbances to dynamic stability, whereas feedforward control relies on knowledge about the expected perturbation generated by prior experience (54, 73). In our study, subjects apparently used feedback control when adapting to the treadmill belt and shift perturbations as MOS and its components did not deviate more from the pre levels with more experience with catch strides. Despite the stride-by-stride exposure to the perturbations, subjects did not appear to develop anticipatory responses. The small perturbations in our study likely contributed to the use of feedback control, as mild medial-lateral perturbations reduce feedforward control and facilitate feedback control in the medial-lateral direction (74). Additionally, the repetitive fixed magnitude treadmill belt perturbations also likely contributed to the use of feedback control. Unlike overground walking where subjects can control the displacement of their stance foot and the magnitude of a slip perturbation which allows for more proactive responses (65), subjects are unable to control the displacement of their stance foot during treadmill belt perturbations, resulting in reactive compensatory step lengths and trunk angles responses (75).

Perturbations in the coincident direction more strongly affected gait stability after being initially perturbed. As one might expect, the anterior-posterior and medial-lateral perturbations resulted in a stronger effect in their respective directions (7, 76). In our study, this directional effect was not merely seen in the MOS_lhs_ but was also reflected in its components. Despite the small magnitude, our perturbations also affected metrics in perpendicular directions, though not as strongly as perturbations in coincident directions. Perturbations in one direction often elicit responses perpendicular to the perturbation direction (4, 7, 76, 77) as responses between the sagittal and frontal planes are dynamically coupled (31, 78). Medial-lateral perturbations are often more challenging than the anterior-posterior perturbations (8, 76) because medial-lateral balance requires active control during walking (47, 79). In our study, medial-lateral perturbations were not more challenging than the anterior-posterior perturbations in the anterior-posterior direction, which could be related to the magnitude of the shifts. We used 1 cm medial-lateral shift perturbations as opposed to the 12-15 cm shifts that were used in other studies (8, 76). This suggests that there is a threshold, a medial-lateral shift > 1 cm that is needed to elicit a significant effect on gait stability compared to small anterior-posterior perturbations. We chose relatively small perturbation magnitudes to avoid falls and extra recovery steps as we applied the perturbations on a stride-by-stride basis.

As expected, there was a scaling effect of perturbation magnitude on the anterior-posterior metrics as people tend to magnify their responses proportionally with perturbation size (7, 26, 77). Changes in MOS_ml, lhs_ and its components were similar between the two treadmill belt perturbations sizes. In our study, the MOS_ml, lhs_ did not respond strongly to the anterior-posterior perturbations (Stick0.4 and Slip0.4) so an even smaller half-sized perturbation (Stick0.2 and Slip0.2) would not be likely to elicit a strong response. Based on the interaction effect for BOS_ap, lhs_ observed in our anterior-posterior perturbations (Fig. 9A), we found that the stick perturbations had a stronger scaling effect than slip perturbations for the same increment (from 0.2 to 0.4). Backward instability created by belt decelerations (stick perturbations) tend to be more challenging than forward instability created by the belt accelerations (slip perturbations) (10, 70).

The calculation of MOS is based on an inverted pendulum model and has several assumptions (19). First, the mass of the whole body is modeled as a single mass point of the inverted pendulum, so the possible effects of upper body (arm or trunk) movements in response to perturbations are ignored (18). Second, the leg is assumed to be rigid and with constant length, which does not consider the large effects of the knee joint motions and moments in response to perturbations (24, 80). Third, the excursions of the COM are small with respect to pendulum length (19), which was likely met in our study due to the small magnitude perturbations. While all of the assumptions were not likely met in our study, the MOS is a common metric of balance for studies of walking with large perturbations (6, 11, 23–26, 29, 30) and the proposed approaches to fulfill the MOS assumptions vary among studies (81).

The present study had several limitations. One limitation of our study is the small sample size. Post-hoc power analyses indicated that hypotheses 1 and 3 were adequately powered while hypotheses 2 and 4 were underpowered, which means we could not reliably detect small statistical differences for all hypotheses. The onsets of the medial-lateral perturbations had a larger delay compared to the anterior-posterior perturbations. The perturbation did not always return to the neutral position or baseline speed before the left heel strike, which likely introduced tiny but extra disturbances. The treadmill system also has a fixed acceleration value for shifting the treadmill platform medial-laterally, which limited the size of the shift perturbation we could use to ≤ 1 cm for our desired time window such that the treadmill could return to its neutral position at or near left heel strike. To control for the time of the perturbation, we set the duration of the anterior-posterior perturbations to be similar to the medial-lateral perturbations, which again limited the magnitude of the perturbation we could apply. We only applied perturbations to the right leg to limit the experiment to ~1 hour. We would not have expected a difference between perturbing the left or right stepping limbs during walking or in forward falls (82, 83). Last, we only applied perturbations at one instance of the gait cycle, right leg mid-stance. A recent study applied brief treadmill belt accelerations at push off of each foot on a stride-by-stride basis and that led to sustained increased push-off once the perturbations were removed (84). Applying perturbations at different points of the gait cycle may reveal phase-dependent modulation of margin of stability and its components.

Future directions include studies to determine the muscular and cortical mechanisms involved during adaptation of gait stability in response to small magnitude perturbations during walking. We recently found significant differences in electrocortical activity of the anterior cingulate and motor cortices in response to small magnitude mechanical perturbations applied on a stride-by-stride basis during recumbent stepping using electroencephalography (EEG) and source estimation (85). Recording EEG and performing source estimation for subjects experiencing small magnitude perturbations on a stride-by-stride basis during walking may reveal specific cortical networks for modulating foot placement versus center of mass when adapting to perturbations during walking. Currently, EEG gait adaptation studies often use large infrequent perturbations during walking (86, 87).

## 5. Conclusions

We investigated how healthy young individuals adjusted their margin of stability, base of support, and extrapolated center of mass in response to small magnitude multidirectional perturbations applied on a stride-by-stride basis. Small magnitude perturbations disrupted gait stability by decreasing the MOS_ap, lhs_ in the belt deceleration (Stick0.4), belt acceleration (Slip0.4), and medial treadmill shift (M1) perturbations, while increasing the MOS_ml, lhs_ in the lateral treadmill shift (L1) perturbation. During adaptation, BOS (foot placement) control was primarily used in the L1 perturbations, whereas XCOM control was preferred in the Slip0.4 and M1 perturbations. Subjects used both feedback and feedforward strategies after being initially perturbed, but primarily used feedback strategies when adjusting their BOS and XCOM during adaptation. Moreover, MOS_lhs_ and its components were generally more sensitive to perturbations in the coincident direction and scaled with perturbation size upon initial exposure to the perturbations. Overall, our findings provide insights into inter-stride gait stability and adaptability to small magnitude perturbations experienced on a stride-by-stride basis and have implications for the development of training interventions that could target specific components of gait stability (e.g., base of support or extrapolated center of mass control) for balance-impaired populations.

## SUPPLEMENTAL DATA

Supplemental Tables S1–S6: https://doi.org/10.6084/m9.figshare.16910683

Supplemental Fig. S1–S2: https://doi.org/10.6084/m9.figshare.16910713

## ACKNOWLEDGMENTS

The authors thank Alejandro Hurtado, Kevin Graydon, Amy Lebanoff, and Allen Rahrooh for assistance with data collection.

## AUTHOR CONTRIBUTIONS

J.L. and H.J.H. conceived and designed research; J.L. performed experiments; J.L. analyzed data; J.L. and H.J.H. interpreted results of experiments; J.L. prepared figures; J.L. drafted manuscript; J.L. and H.J.H. edited and revised manuscript; J.L. and H.J.H. approved final version of manuscript.

## ENDNOTE

At the request of the authors, readers are herein alerted to the fact that additional materials related to this manuscript may be found at https://doi.org/10.6084/m9.figshare.c.5285258.v3. These materials are not a part of this manuscript and have not undergone peer review by the American Physiological Society (APS). APS and the journal editors take no responsibility for these materials, for the website address, or for any links to or from it.

---

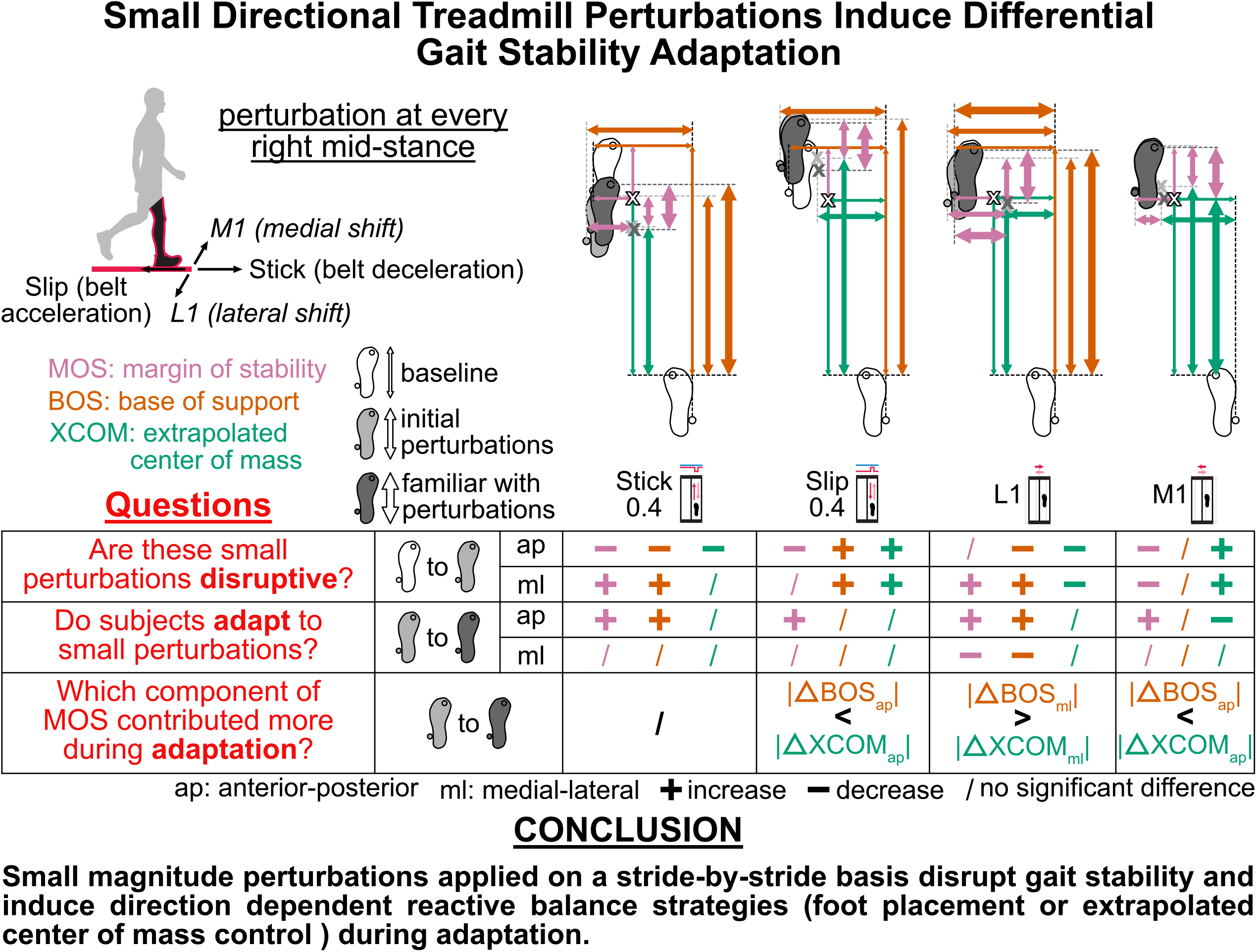

